# The variation landscape of *CYP2D6* in a multi-ethnic Asian population

**DOI:** 10.1101/2024.01.20.576401

**Authors:** Yusuf Maulana, Rodrigo Toro Jimenez, David Twesigomwe, Levana Sani, Astrid Irwanto, Nicolas Bertin, Mar Gonzalez-Porta

## Abstract

Cytochrome P450 2D6 (CYP2D6) plays a crucial role in metabolizing approximately 20% of medications prescribed clinically. This enzyme is encoded by the *CYP2D6* gene, known for its extensive polymorphism with over 170 catalogued haplotypes or star alleles, which can have a profound impact on drug efficacy and safety. Despite its importance, a gap exists in the global genomic databases, which are predominantly representative of European ancestries, thereby limiting comprehensive knowledge of *CYP2D6* variation in ethnically diverse populations. In an effort to bridge this knowledge gap, we focused on elucidating the *CYP2D6* variation landscape within a multi-ethnic Asian cohort, encompassing individuals of Chinese, Malay, and Indian descent. Our study comprised data analysis of 1,850 whole genomes from the SG10K_Health dataset using an in-house consensus algorithm, which integrates the capabilities of Cyrius, Aldy, and StellarPGx. This analysis unveiled distinct population-specific star-allele distribution trends, highlighting the unique genetic makeup of the Singaporean population. Significantly, 46% of our cohort harbored actionable *CYP2D6* variants—those with direct implications for drug dosing and treatment strategies. Furthermore, we identified 14 potential novel *CYP2D6* star-alleles, of which 7 were observed in multiple individuals, suggesting their broader relevance. Overall, our study contributes novel data on *CYP2D6* genetic variations specific to the Southeast Asian context. The findings are instrumental for the advancement of pharmacogenomics and personalized medicine, not only in Southeast Asia but also in other regions with comparable genetic diversity.

## Introduction

Pharmacogenomics entails understanding how genetic variation affects individual responses to drugs and it stands as a cornerstone in the advancement of personalized medicine. Within the landscape of drug-metabolizing enzymes, cytochrome P450 2D6, encoded by the *CYP2D6* gene, is well-recognized as a pivotal player. While it represents only 2-4% of the total CYP enzymes in the liver, CYP2D6 is responsible for metabolizing approximately 20% of clinically prescribed drugs, including a wide array of medications such as beta blockers, antiarrhythmics, antidepressants, antipsychotics, anticancer agents like tamoxifen, and opioids^1^. The significance of *CYP2D6* in pharmacogenomics is further emphasized by its extensive genetic diversity. To date, the Pharmacogene Variation (PharmVar) Consortium catalogue contains over 170 *CYP2D6* haplotypes, commonly referred to as star alleles^2,3^. The majority of these star alleles are defined by specific combinations of single nucleotide polymorphisms (SNPs) and/or small insertions and deletions (indels). However, known variations in the *CYP2D6* locus also encompass complex structural variants (SVs), such as full gene deletions, duplications, multiplications, and hybrid tandem rearrangements involving the closely related *CYP2D7* and *CYP2D8* pseudogenes.

The vast genetic heterogeneity of *CYP2D6* profoundly influences the effectiveness and safety of drugs metabolized by the CYP2D6 enzyme^1^. For instance, poor metabolizers carry two non-functional alleles and may not effectively metabolize or bioactivate drugs through the CYP2D6 pathway. Conversely, ultra-rapid metabolizers, who possess at least one increased function allele in addition to a normal-function allele, are at heightened risk of experiencing dose-related adverse events or treatment ineffectiveness. Thus, genetic variations can serve as pharmacogenetic biomarkers to guide drug dosing decisions, improve the effectiveness of therapies, and prevent adverse reactions for drugs metabolized by CYP2D6^4^.

In this context, the Clinical Pharmacogenetics Implementation Consortium (CPIC) has developed guidelines to aid in translating pharmacogenetic laboratory test results into actionable prescribing decisions for specific drugs, including three dedicated to CYP2D6^5^. However, the frequency of *CYP2D6* alleles exhibits considerable variability across different global populations, with some allelic variants universally present and others varying significantly in frequency or being exclusive to certain ethnic groups^4,6–9^. This diversity highlights the critical need to understand the variation landscape of *CYP2D6* for effective pharmacogenomic applications, particularly in the context of large-scale population genomic studies that can help inform drug policies. Nonetheless, there remains a noticeable gap in the data concerning variant frequencies in pharmacogenes, especially within Asian populations, since the majority of genetic variation data in public databases primarily originate from Western populations.

Singapore, with its diverse ethnic population, presents a unique opportunity for inclusive and comprehensive pharmacogenomic research. In addition, the advent of next-generation sequencing (NGS) technologies has made it feasible to sequence large cohorts, and with the aid of dedicated callers, NGS has demonstrated its capability to accurately characterize haplotypes in the *CYP2D6* gene, even in the presence of the aforementioned complexities^10^. In this context, our study aims to leverage Singapore’s highly diverse genetic landscape, coupled with the capabilities of whole genome short-read sequencing, to provide insights into the variation landscape of the *CYP2D6* gene within Asian groups. To achieve this, we introduce a newly developed multi-tool bioinformatics workflow, and we apply it to a unique cohort comprising over 1,400 samples that have been sequenced at high depth using short reads^11^. We use the results of these analyses to investigate the distribution of known and potential novel *CYP2D6* star alleles in the three majority Asian ethnicities in Singapore (Chinese, Indian and Malay), and to characterize the prevalence of different metabolizer profiles in the population. The insights garnered from this research hold the potential for significant implications in the realm of clinical pharmacogenetics implementation strategies, not only within Singapore but also across Southeast Asia.

## Results

### Development and evaluation of a multi-tool pipeline for *CYP2D6* star allele calling

We developed a multi-tool analysis pipeline (**Figure 1A**) to resolve *CYP2D6* diplotypes using three publicly available callers: Cyrius^12^, Aldy^13^, and StellarPGx^14^. Our pipeline, implemented in Nextflow (see **Methods**), accepts whole genome sequencing Compressed Reference-oriented Alignment Map (CRAM) files as input. It initially generates individual calls from each of the three callers, which are then subjected to a consensus algorithm. This algorithm reports a consensus call when at least two out of the three callers concur on the diplotype. Typically, ambiguous calls are assigned for samples where this consensus criteria is not met. However, we manually inspected cases where the presence of potential novel alleles had been predicted by StellarPGx to further characterize and report the novel core variant combinations.

**Figure 1:**
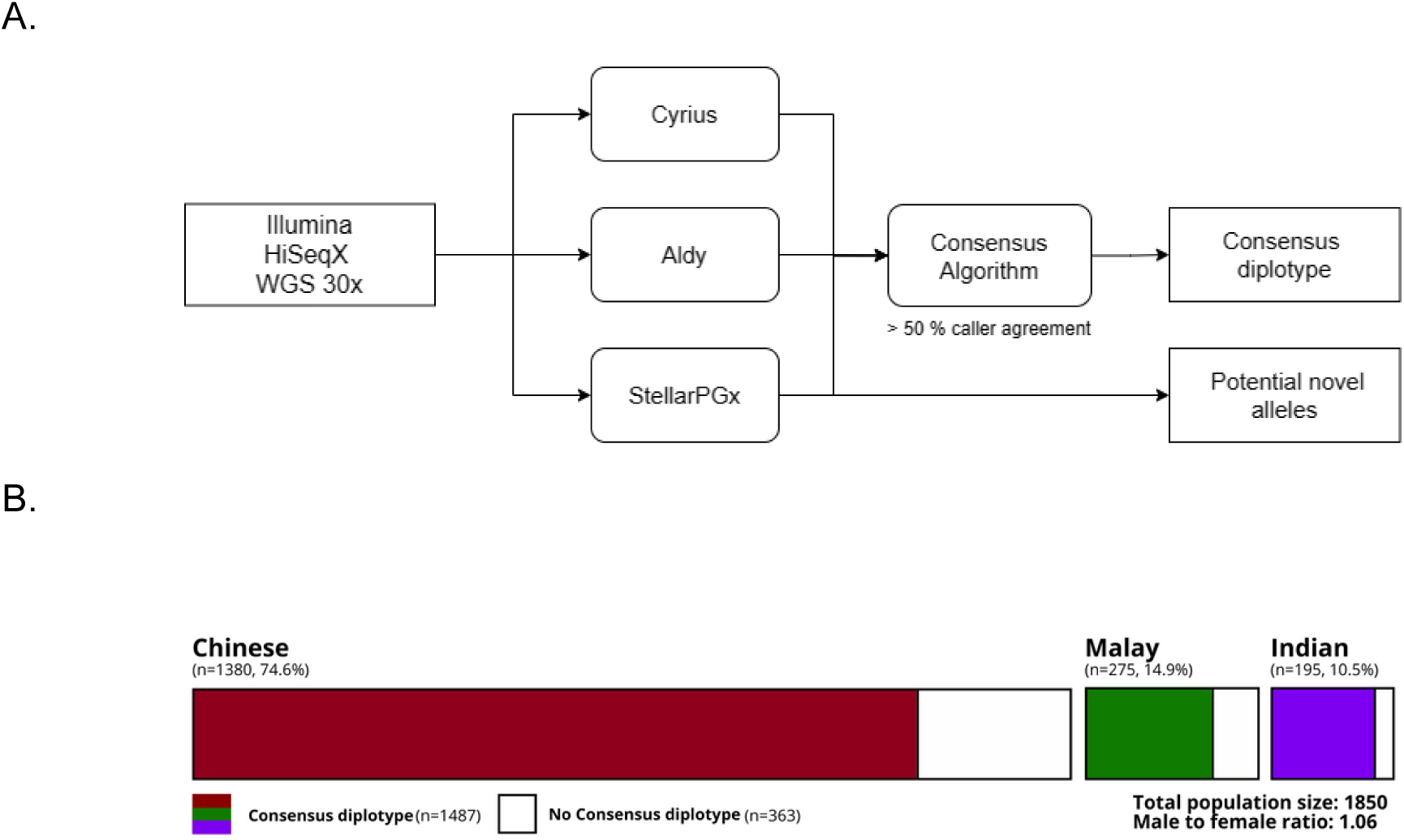
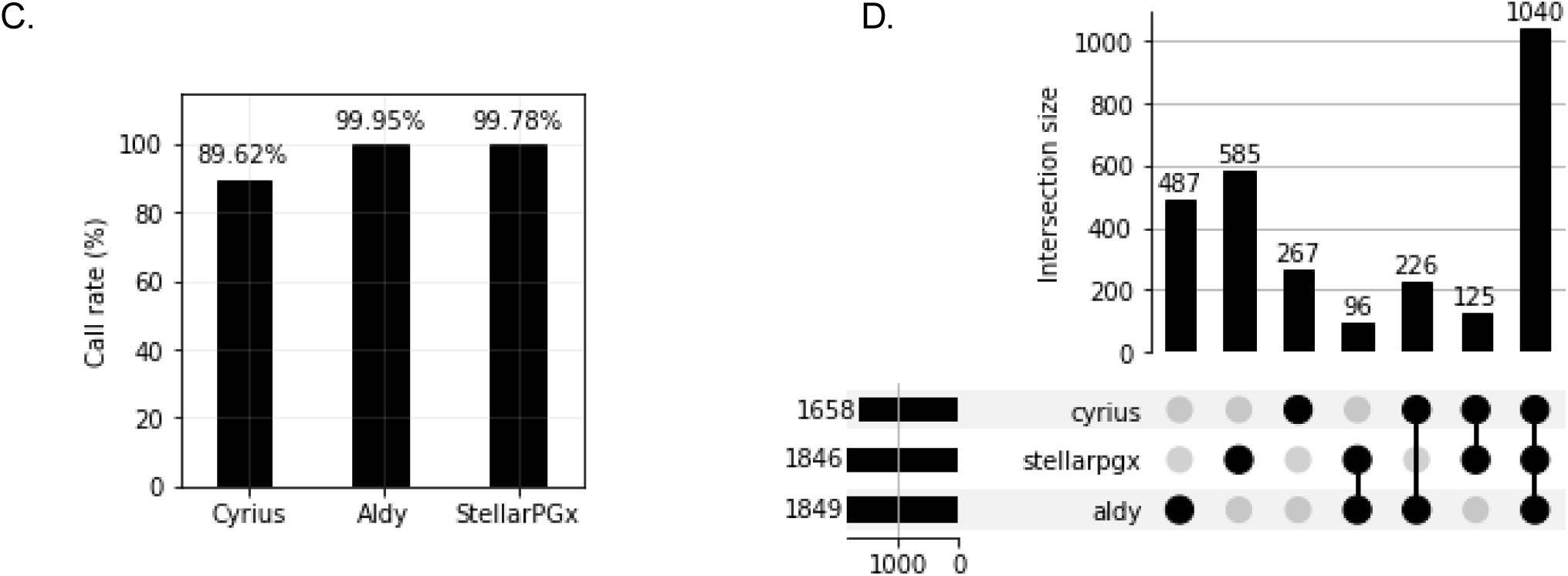
Overview of data analysis workflow and study design. **A. Data analysis pipeline.** The diagram illustrates the workflow for analyzing WGS 30X CRAM files. Three tools —Cyrius, Aldy, and StellarPGx— are utilized for *CYP2D6* genotyping. A consensus algorithm determines the final diplotypes based on agreement from at least two of the three tools. StellarPGx is employed to identify potential novel alleles. **B. SG10K_Health cohort.** The cohort for the study comprises individuals from the three major ethnic groups in Singapore, namely Chinese, Malay, and Indian. Colored bars represent the count and proportion of consensus diplotypes within the total cohort of 1,850 participants. **C. Comparison of call rates**. The bar graph compares the successful calling rates of Cyrius, Aldy, and StellarPGx in generating diplotype calls. **D. UpSet plot of CYP2D6 diplotype predictions.** The plot displays the overlap in diplotype calls among the three callers. Numbers atop the bars show the size of each intersection set. The lower left side of the plot indicates the total successful calls per tool. The unique diplotypes identified by each tool are shown in the three leftmost bars (Aldy with 487, StellarPGx with 585 and Cyrius with 267 unique diplotype calls), the middle bars represent diplotypes identified by tool pairs, and the far-right bar shows 1,040 diplotypes identified by all three tools.

To characterize the variation landscape of *CYP2D6* in a multi-ethnic population, we executed our analysis pipeline on a dataset comprising 1,850 samples from SG10K_Health, consisting of unrelated and healthy study participants (**Figure 1B**). This dataset represents a highly diverse Southeast Asian cohort, including individuals of Chinese (74.6%), Malay (14.9%), and Indian (10.5%) ancestries, and was generated via short-read Whole Genome Sequencing (WGS) at 20-30x coverage. We first examined the call rates of each individual caller and observed that Aldy and StellarPGx exhibited the highest call rates at 99.95% and 99.78%, respectively, while Cyrius employed a more conservative approach with a call rate of 89.62% (**Figure 1C**). Next, we assessed the concordance of diplotype calls across the three tools and found that 56.59% (N=1,040) of calls were supported by all three callers, 80.37% (N=1,487) were supported by at least two callers, and 19.62% (N=363) were either supported by only one caller or harbor potential novel alleles. In pairwise comparisons, Cyrius and Aldy demonstrated the highest level of agreement, sharing 1,266 haplotype calls (including 1,040 shared among the three callers), followed by StellarPGx and Cyrius with 1,165 shared calls, and StellarPGx and Aldy with 1,136 shared calls (**Figure 1D**). Notably, StellarPGx exhibited the ability to resolve the greatest number of unique diplotypes (N=585), followed by Aldy (N=487) and Cyrius (N=267), which could be attributed to StellarPGx’s capacity to detect novel alleles. In total, we identified 93 samples that potentially contained novel alleles, representing approximately 5% of our study population, which were further refined to a smaller set of 28 samples (1.5%) following manual curation (see below).

### *CYP2D6* star allele frequencies and correlation with PharmGKB

In our subsequent analysis, we aimed to characterize the prevalence of star alleles within the different ancestral groups in our study cohort. **Figure 2A** presents the frequencies of the ten most common haplotypes, categorized by genetic ancestry, and reveals distinct patterns in star allele frequencies across the various ethnic groups. For example, the Chinese and Malay populations predominantly exhibit the \**36+*10*, *\*10*, and *\*36* alleles. In contrast, the Indian population displays a higher frequency of the *\*2*, *\*41*, *\*5*, and *\*4* alleles. Notably, 80% of the top ten most prevalent haplotypes contain actionable variants, characterized as either non-functional or reduced function *CYP2D6* alleles. In addition, we have detected 11 haplotypes that are only present in one individual (*\*112, *52, *133, *7, *75, *82, *17, *9, *15, *4+*4, *69*). (**Supplementary Table 1**).

**Figure 2:**
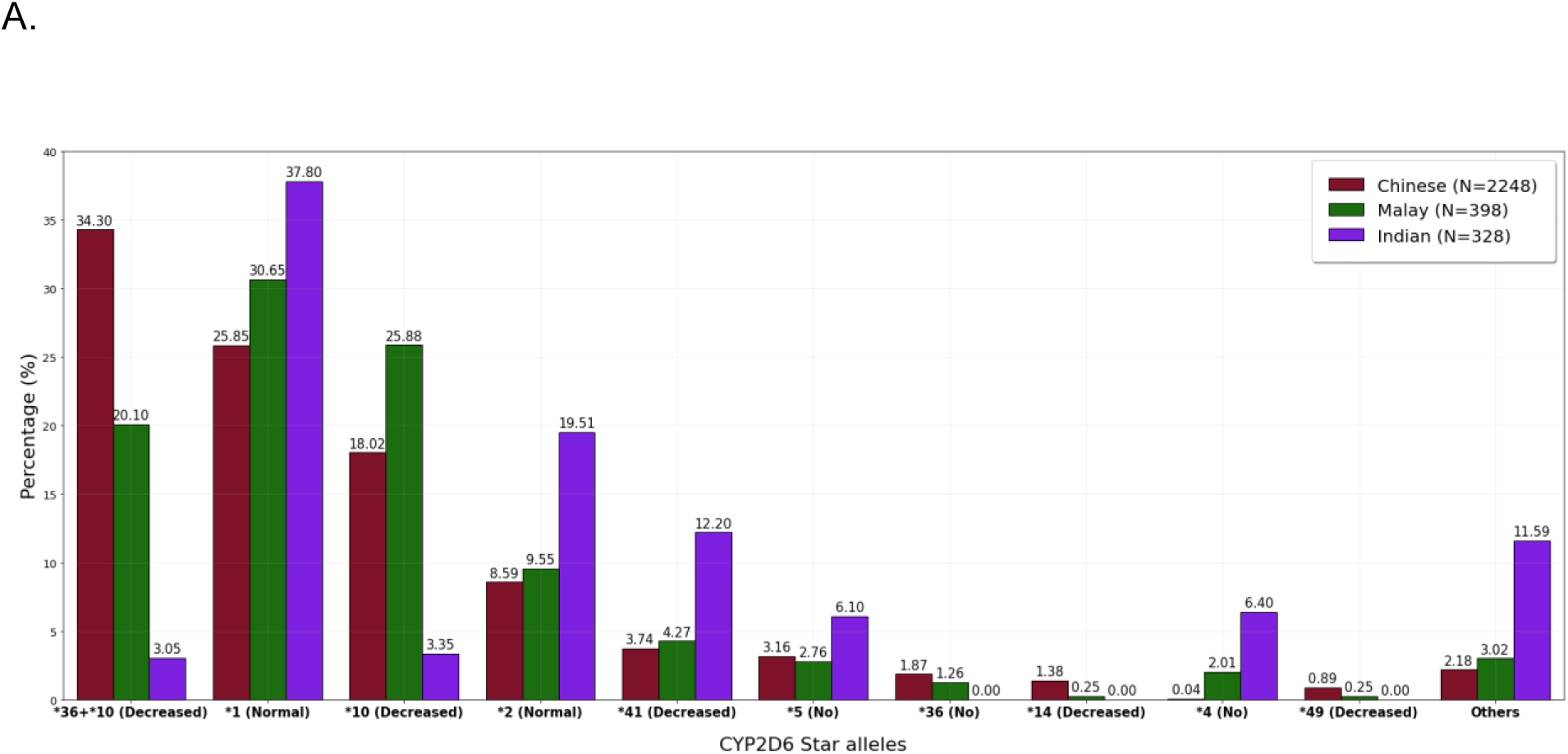
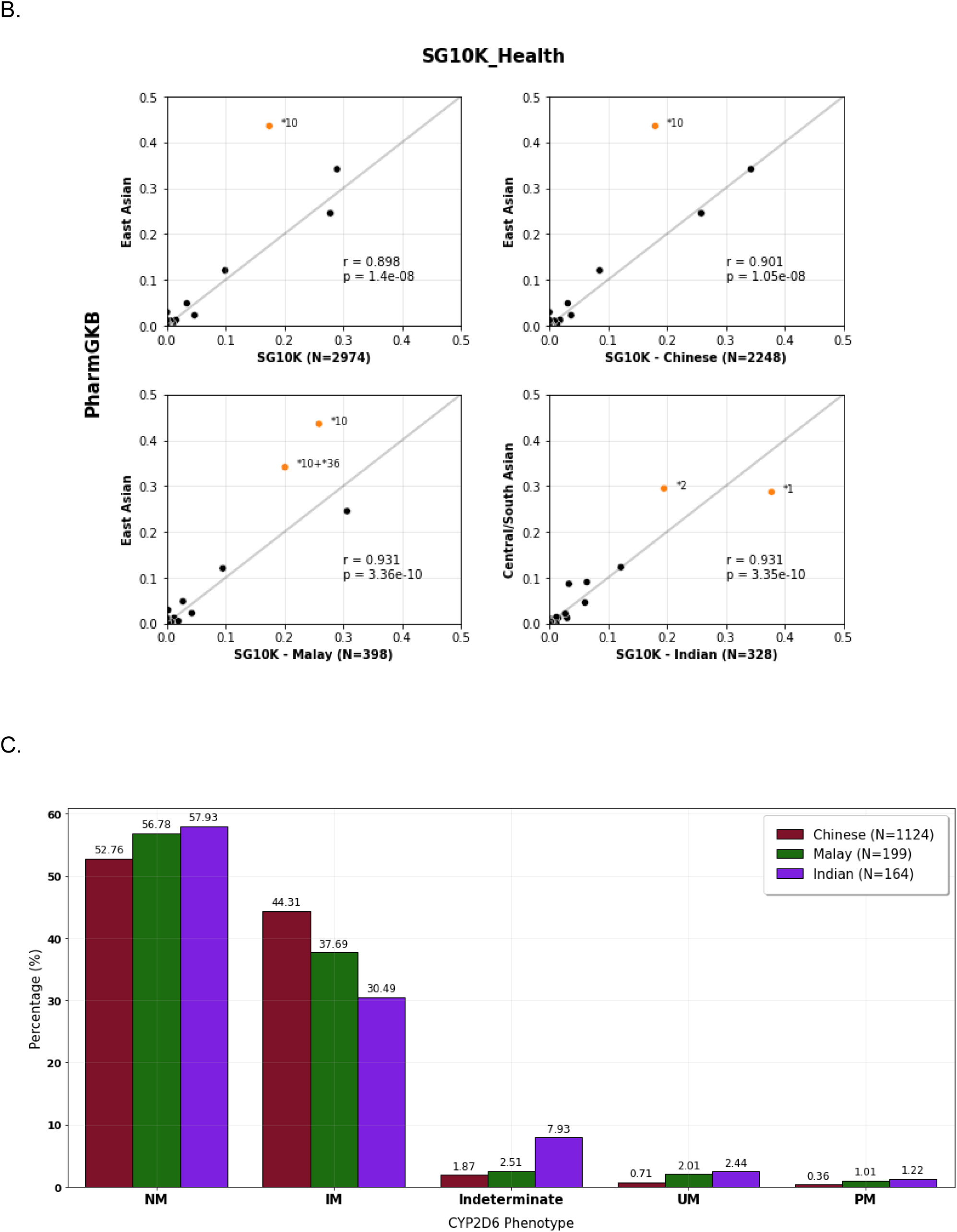
Distribution of *CYP2D6* star alleles and phenotypes across different genetic ancestries. **A. Frequency distribution of the top 10 *CYP2D6* star alleles in the SG10K_Health dataset, categorized by genetic ancestries.** The x-axis depicts the star alleles, each annotated with their respective functional classification (normal, decreased, or no function). The y-axis shows the frequency distribution (%) of these star alleles within each genetic ancestry group. **B. Comparative analysis of *CYP2D6* star allele frequencies between the SG10K_Health dataset and PharmGKB.** The x-axis displays the frequency of star alleles observed in the SG10K_Health dataset. The y-axis indicates the frequency of star alleles as reported in PharmGKB for East Asian and Central/South Asian populations. The Pearson correlation coefficient (*r*) is represented for each comparison. **C. Frequency distribution of CYP2D6 phenotypes among different genetic ancestries.** The x-axis lists all the possible CYP2D6 phenotypes, including normal metabolizers (NM), intermediate metabolizers (IM), indeterminate, ultrarapid metabolizers (UM), and poor metabolizers (PM). The y-axis shows the frequency distribution (%) of these phenotypes across each genetic ancestry.

Furthermore, we conducted a comparison of haplotype frequencies between the SG10K_Health and PharmGKB datasets, revealing a strong correlation with coefficients surpassing 0.90 (**Figure 2B**). Importantly, this trend remains consistent across individual ethnicities. However, we noted a discrepancy in the frequency of the *\*10* and *\*36+*10* alleles in Chinese and Malay ethnicities, both lower in the SG10K_Health dataset compared to the East Asian population in PharmGKB.

### Distribution of CYP2D6 Metabolizer Profiles

Among the 1487 samples that met the criteria of our consensus algorithm, approximately 46.14% exhibited actionable CYP2D6 phenotypes affecting drug metabolism (i.e., intermediate metabolizer (IM), poor metabolizer (PM), and ultrarapid metabolizer (UM) phenotypes), emphasizing the role that pharmacogenomics can play in clinical practice. Further analysis of phenotype distribution in our cohort revealed that 53.87% of the study samples could be classified as normal metabolizer (NM), followed by 41.90% as IM, then 1.08% as UM, and 0.54% as PM (**Supplementary Figure 1**). IMs and PMs could be attributed to the higher prevalence of alleles and nonfunctional alleles such as *\*10*, *\*36*, *\*41* and *\*5*. Additionally, 2.62% of the samples had indeterminate phenotypes, representing variants with unknown effects on metabolism. We also identified three instances of complete *CYP2D6* deletions (\**5/*5*).

Our study also unveiled distinct patterns of metabolizer profiles within the SG10K_Health cohort based on ancestry (**Figure 2C**). Notably, a lower incidence of NM was observed in the Chinese and Malay participants, possibly linked to the higher prevalence of *\*36+*10* hybrids. In contrast, individuals of Indian descent exhibited a higher prevalence of indeterminate phenotypes (7.93% compared to 2.51% and 1.87% for Malay and Chinese, respectively), associated with the prevalence of star alleles with unknown function (\**43*, *\*86*, *\*113*, *\*82*, *\*111*, and *\*112*).

### Distribution of *CYP2D6* Structural Variants

Subsequently, we aimed to characterize *CYP2D6* Structural Variants (SVs) within the SG10K_Health cohort. As expected, the most prominent SV-containing star allele was *\*36+*10*, which displayed an average frequency of 28.95% in the total population, primarily contributed by Chinese and Malays (**Figure 3A**). This was followed by *CYP2D6*5* (full-gene deletion), albeit at a much lower frequency of 3.43% overall, and with a two-fold higher prevalence in Indians. Additionally, we also detected *\*2×2* and *\*1×2* haplotypes, indicative of gene duplications, with frequencies of 0.34% and 0.3% across the total population, respectively. These haplotypes showed increased function due to the multiplication of normal function alleles (*\*2* and *\*1*). Lastly, the *\*68+*4* hybrid and *\*4+*4* allele, both characterized by having no function, were less common in the overall cohort (0.37%) and were predominantly found in the Indian subgroup.

**Figure 3:**
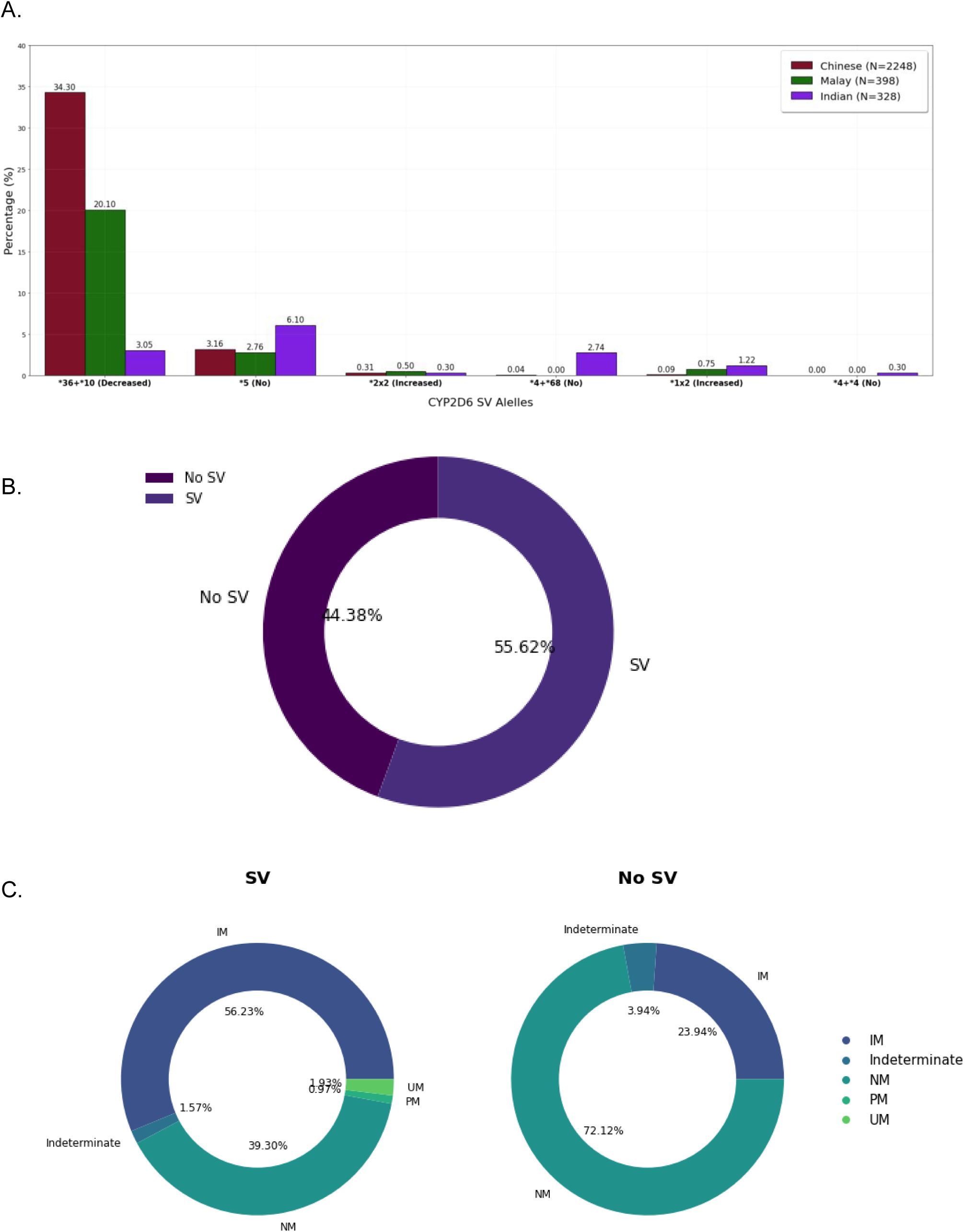
Frequency distribution of *CYP2D6* structural variations (SVs). **A. Frequency distribution of SV-containing *CYP2D6* star alleles in the SG10K_Health dataset, categorized by genetic ancestries.** The x-axis depicts the star alleles, each annotated with their respective functional classification (increased, decreased, or no function). The y-axis shows the frequency distribution (%) of these star alleles within each genetic ancestry group. **B. Comparison of the proportion of samples with SV and samples with no SV in the SG10K_Health dataset (N=1487). C. Distribution of CYP2D6 phenotypes in the SG10K_Health dataset, categorized by SV presence.** The left side illustrates the proportion in samples with SV (N=827), and the right side shows the proportion in samples with no SV (N=660).

Notably, the majority of participants (55.62%) harbored at least one star allele containing an SV, while 44.38% harbored no observable SVs (**Figure 3B**). However, these results may constitute an under-representation of the prevalence of SVs in our study population, given the stringency of the consensus algorithm that we employed and the inherent limitations of short-reads. Stratification based on the presence or absence of SVs revealed significant differences in metabolizer profiles (**Figure 3C**). As expected, in samples without SVs, NM made up 72.12% of the cases, followed by IM at 23.94%, and a smaller proportion of indeterminate samples at 3.94%. In contrast, the group with SVs predominantly consisted of IM (56.23%). NM constituted the second most common category, with 39.30% frequency, highlighting the importance of detecting the exact nature of the *CYP2D6* SVs present in a given sample. We also identified UM and PM, though they were less prevalent, with frequencies of 1.93% and 0.97%, respectively.

### Potentially Novel *CYP2D6* Alleles

Lastly, we proceeded to examine the set of potentially novel star alleles detected by StellarPGx. Initially, this tool flagged 93 samples as containing potential novel alleles; however, following manual review, we refined the dataset to 13 computationally-resolved potential novel haplotypes and 1 potential novel allele in a diplotype with unresolved phasing (**Table 1** and **Figure 4**). The remaining samples did not exhibit sufficient evidence to be considered as harboring novel alleles, with the majority presenting no additional core variants after manual curation (N=37) while the rest are characterized by low quality or potential miscalls indicative of existing combination of star alleles. Among the 14 potential novel alleles, the majority were associated with the *\*10* and *\*2* haplotype backbones (9 out of 14). Notably, these novel alleles were observed in 28 study samples, with 7 of these alleles detected in multiple individuals, and 4 samples from different ethnic backgrounds, consistently observed in both Chinese and Malay individuals. In contrast, among single-population occurrences, 7 unique events were exclusive to the Chinese group, while 2 were unique to the Indian group. All detected novel alleles in our study were found to contain functional variants, including missense and frameshift mutations, which potentially alter protein functionality. While these novel core variants are cataloged in the dbSNP database, they are novel in the context of our study as they appear in combinations not previously observed with *CYP2D6* alleles in the PharmVar database. Noteworthy, 4 of the allele-defining variants— rs746803316, rs3915951, rs1135830, and rs759234339— have not been previously catalogued as *CYP2D6* core variants in the PharmVar database.

**Figure 4:**
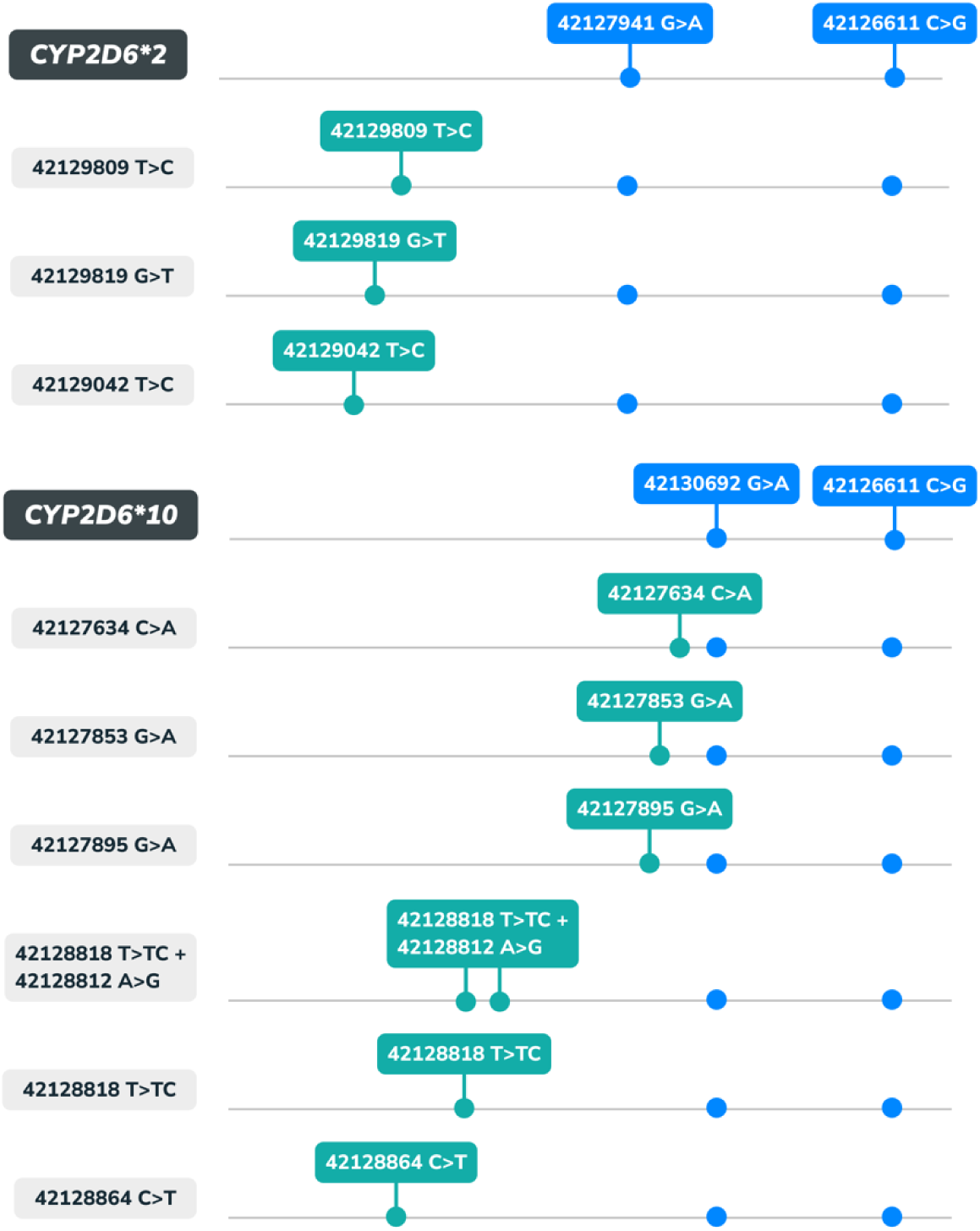
Visual representation of potential novel alleles with *\*2* and *\*10* background alleles. The core variants of the background alleles are marked in blue, and the unique novel variants are marked in green.

**Table 1:** Potentially novel *CYP2D6* star alleles. The 14 potentially novel haplotypes identified following manual curation of StellarPGx outputs.

| Haplotype | Background allele | Additional core variant(s) |  | Variant type | AF gnomAD |  | Count |
| --- | --- | --- | --- | --- | --- | --- | --- |
|  |  | GRCh38 (NC_000022.11) | Rsid |  | Global | East Asian |  |
| 1 | *1 | 42128341C>T | rs746803316 | Missense(A226T)<sup>†</sup> | 2.73.E-05 | 8.27.E-04 | 3 |
| 2 | *10 | 42127634C>A | rs3915951 | Missense(R329L)<sup>†</sup> | 2.66.E-03 | 7.56.E-03 | 6 |
| 3 | *10 | 42127853G>A | rs140513104 | Missense(P325L) | 1.53.E-04 | 4.69.E-04 | 4 |
| 4 | *10 | 42128812A>G,<br>42128818T>TC | rs199535154,<br>rs72549354 | Missense(L213P),<br>Frameshift(L213fs) | 2.54.E-04, 0 | 5.64.E-04, 0 | 2 |
| 5 | *10 | 42127895G>A | rs1135830 | Missense(S311L)<sup>†</sup> | 6.77.E-05 | 6.71.E-05 | 2 |
| 6 | *10 | 42128864C>T | rs755518310 | Missense(E196K) | 8.11.E-06 | 2.25.E-05 | 1 |
| 7 | *10 | 42128818T>TC | rs72549354 | Frameshift(L213fs) | 0 | 0 | 1 |
| 8 | *2 | 42129042T>C | rs1135824 | Missense(N166D) | 1.58.E-04 | 1.79.E-04 | 2 |
| 9 | *2 | 42129819G>T | rs28371703 | Missense(L91M) | 1.50.E-01 | 8.06.E-04 | 2 |
| 10 | *2 | 42129809T>C | rs28371704 | Missense(H94R) | 1.59.E-01 | 7.39.E-04 | 1 |
| 11 | *41 | 42126747G>A | rs730882251 | Missense(R441C) | 1.21.E-04 | 0 | 1 |
| 12 | *86 | 42130715C>T | rs28371696 | Missense(R26H) | 1.52.E-03 | 3.15.E-04 | 1 |
| 13 | *86 | 42130692G>A | rs1065852 | Missense(P34S) | 2.12.E-01 | 4.88.E-01 | 1 |
| **Total** |  |  |  |  |  |  | **27** |
| **Unresolved diplotypes with potentially novel star alleles** |  |  |  |  |  |  |  |
| 1 | *1/*2 | 42129807C>T | rs759234339 | Missense(G283S)<sup>†</sup> | 8.72.E-06 | 0 | 1 |
| **Total** |  |  |  |  |  |  | **1** |

<sup>†</sup> Variants not currently listed in PharmGKB, for which interpretation was conducted using VEP.
AF, Allele Frequency

## Discussion

While extensive research has been conducted on *CYP2D6* across various ethnicities, a gap still exists in understanding the extent of *CYP2D6* pharmacogenetic diversity within Southeast Asian populations. To address this gap, we conducted a study to characterize the distribution of *CYP2D6* star alleles and their associated phenotypes using a genetically diverse cohort from Singapore^11^. This cohort comprises individuals representing the three major ethnicities in the country: Chinese, Malay, and Indian, and includes high-coverage short-read whole genome sequences from over 1,800 participants. To the best of our knowledge, this study represents the most comprehensive examination of *CYP2D6* genetic variation in the Singaporean population, and given the country’s rich diversity, it provides an ideal platform for comprehensively exploring *CYP2D6* variation within Southeast Asia.

We developed a bioinformatics workflow using three distinct tools to mitigate inaccuracies in identifying *CYP2D6* variants and diplotypes, given the limitations presented by the short-read data we used in this study. Our workflow includes a consensus algorithm, which reports a diplotype call only when at least two out of three tools concur; however, cases where potential novel star alleles were predicted by StellarPGx were subjected to additional manual inspection to ascertain the final diplotype calls. This approach, while conservative, was chosen to prioritize the accuracy of star allele assignments. Additionally, our workflow includes steps to interpret predicted diplotypes into metabolizer profiles and to identify potential novel alleles. Upon applying the workflow to the 1,850 samples in our cohort, we successfully determined consensus diplotype calls for 1,487 samples, encompassing over 80% of the population. In contrast, around 20% of samples remained uncharacterized. Notably, the majority of samples that did not reach consensus contained haplotypes that included SVs and potential novel alleles (90.39%), thus underscoring the challenges associated with the genotyping of the *CYP2D6* locus via short-read sequencing. Such challenges arise from the gene’s significant homology with the *CYP2D7* and *CYP2D8* pseudogenes, hinting at the potential for future research utilizing novel technologies such as long-read sequencing.

In our study, we observed significant patterns in the distribution of *CYP2D6* alleles among the Southeast Asian populations we examined. Among the most prevalent star alleles, we noted a predominance of alleles associated with reduced or absent function, except for *\*1* and *\*2*, which are associated with normal function. *\*1* emerged as the most prevalent *CYP2D6* allele with normal function, followed by *\*2*, aligning with previously reported trends in Southeast Asian populations^9^. Additionally, we observed variations in allele frequencies among the Singaporean populations included in our study. *\*36+*10*, *\*10*, and *\*36* were more prevalent in the Chinese and Malay populations, with the Chinese and Malays exhibiting approximately six-fold higher allele frequencies of *\*10* compared to Indians. This trend aligns with well-documented findings, which consistently report a high prevalence of the *\*36+*10* tandem in East Asian populations, including Japanese, Korean, and Chinese^8,9,16^. Furthermore, it supports the common identification of the *\*10* allele as a reduced-function variant in Asian populations^8,9^. Notably, *\*36* was not observed in the Indian participants in this study. In contrast, *\*2*, *\*41*, *\*5*, and *\*4* alleles exhibited higher prevalence in the Indian population, reaffirming previous research highlighting the prominence of these alleles in Indian samples^15^. We also observed 11 haplotypes (*\*112, *52, *133, *7, *75, *82, *17, *9, *15, *4+*4, *69*) that are only present in one individual which expands the previous findings of rare stare alleles in our study population^15^. Of particular interest was the identification of three individuals with no detectable copies of *CYP2D6* (*\*5/*5\**), constituting approximately 0.2% of our population. This aligns with previous reports indicating that the *\*5/*5* diplotype occurs at a very low frequency, ranging from 0% to 1.86% in Southeast Asian populations^9^.

The frequencies of common star alleles in our population, such as *\*10, *36+*10,* and *\*2*, exhibited significant differences from the average frequencies estimated by the PharmGKB and 1000 Genomes Project (1KGP) for the equivalent populations. Whilst PharmGKB may include additional ethnicities to the ones in SG10K_Health, these discrepancies may originate from variations in data generation methods (e.g., WGS versus genotyping) and star allele calling approaches employed in various studies, leading to variability in the range of detectable star alleles. It is also possible that the frequency of *\*10* could be overestimated when *\*36+*10* tandems and/or *\*36* alleles are not reported. Additionally, *\*2* alleles are considered “backbone” alleles since their defining SNVs occur in multiple other haplotypes, which may introduce potential mis-assignments if any of these additional haplotype-defining variants cannot be accurately detected.

After translating diplotypes into metabolizer profiles, we identified actionable variants in over 46% of the population, increasing to over 80% when focusing on the top ten most common haplotypes. This underscores the significant impact of implementing pharmacogenomics on a large scale. In our study population, normal metabolizers (NMs) were the most prevalent phenotype at 53.87%, ultra-rapid metabolizers (UMs) accounted for 1.08%, while poor metabolizers (PMs) represented 0.54%. The highest frequencies of diplotypes predicting PM were found in Indian subjects (1.22%), followed by Malays (1.01%) and Chinese (0.36%). Interestingly, we detected PMs in all three ethnicities (Chinese, Indian, and Malay), a difference from previous studies in the same population that did not identify PMs in Chinese individuals^15^, likely due to our larger sample size and higher-depth sequencing approach. Compared to global trends, our cohort showed a slightly higher incidence of intermediate metabolizers, around 42%, surpassing the previously reported 34%^17^. This coincided with a decrease in the prevalence of normal metabolizers, typically observed at 64-68% in global populations. This trend may be attributed to the higher prevalence of diplotypes associated with reduced or no function, including *\*36+*10, *10, *41*, and *\*36*.

Both PMs and UMs exhibit altered capacity to metabolize CYP2D6 substrates, including codeine, certain antidepressants, and antipsychotics^18^. UMs face an elevated risk of toxicity due to increased morphine formation after codeine administration, while individuals with non-functional alleles are at risk of inadequate pain relief due to reduced efficacy. In our study, the proportion of UMs exceeded that of PMs by approximately than two-fold. Unlike Caucasians, where the *\*4* allele predominates and accounts for 70-90% of poor metabolizer status, its low frequency in Asians may explain the lower proportion of poor metabolizers in our population^17^. Lastly, we observed a relatively high proportion of Indian participants (7.93%) with an indeterminate CYP2D6 metabolizer phenotype, highlighting the limitations of current CPIC guidelines for genotype-phenotype translation. Most of these individuals carried alleles with uncertain functions, such as *\*43, *86, *113, *82, *111*, and *\*112*. This underscores the need for extensive allele characterization and phenotypic studies to develop effective precision medicine strategies, particularly for medications metabolized by CYP2D6.

We further inspected the prevalence of SVs in *CYP2D6*, as it remains unexplored in Asian populations^8^. Our analysis indicated that the majority of study participants (55.62%) had at least one SV-containing star allele, with *\*36+*10* hybrids being the most prevalent overall. This percentage significantly exceeds previously reported rates, and it is likely to constitute an underestimation of the true prevalence of SVs, given the limitations of short-read sequencing in uncovering such type of variation^19^. When stratifying metabolizer profiles based on the presence or absence of SVs, we detected a higher incidence of IMs (56.23%), UMs (1.93%) and PMs (0.97%) among participants with SV-containing alleles compared to individuals with no *CYP2D6* SVs, as expected. Interestingly, we still detected a high prevalence of NMs among the first group (39.30%), highlighting the importance of detecting the exact nature of the *CYP2D6* SVs present in each sample, and emphasizing the limitations of relying on copy number or SV information alone to make predictions on phenotypic outcomes.

Lastly, our study also provides an initial assessment of the extent of genetic variation that remains undocumented in public databases. We identified 14 potential novel alleles for *CYP2D6*, based on a carefully curated subset. This group includes both shared variants observed across multiple individuals (N=7) and private events (N=7). Although these novel star alleles are individually rare, collectively, they appear in 28 individuals, accounting for 1.5% of our study population. This underscores the significant, yet often overlooked, impact that rare allelic variations could have on precision medicine strategies, both in Asia and globally. The majority of these 13 novel alleles are variations of the *\*10* allele (N=6), followed by *\*2* (N=3). This distribution aligns with the prevalence of these alleles, which are among the top ten star alleles frequently identified in our study populations. All the novel alleles we detected include potentially functional variants previously catalogued in PharmVar in combination with other haplotypes, except for 5, which we inferred using the Variant Effect Predictor. They encompass a range of genetic changes, including missense and frameshift mutations, which could significantly alter protein function. However, given the complexities of genotyping and star allele calling in *CYP2D6*, caution is advised when interpreting computationally inferred novel alleles, particularly singletons. The predictions can be influenced by limitations in the genomic datasets used, such as short-read sequencing and software. While these computational predictions are informative, they are not a substitute for actual experimental validation. Although we could not validate these alleles due to lack of DNA access, our findings offer a valuable foundation for future research, especially as Singapore’s National Precision Medicine program progresses in characterizing a larger portion of its population. A prevalent practice in the field is to report variants based on their backbone allele, which could significantly influence drug dosage recommendations. Unexamined variants with potential functional significance, like those uncovered in our study, could affect phenotype assignments and clinical decisions, potentially leading to adverse pharmacological outcomes for patients.

Overall, our research represents a significant step towards enhancing the understanding of *CYP2D6* variations in under-represented populations. Gaining a thorough insight into the allelic diversity within the populations in this study is key for precisely predicting drug responses and successfully applying pharmacogenetics in Singaporean and global clinical settings, where targeted assays are still the norm. Therefore, understanding the genetic variability of the various populations is essential to ensure that the most prevalent alleles are effectively identified and included in the list of targets. While a broader test such as whole genome sequencing would be ideal to continuously explore genetic variation and guide future clinical applications, cost constraints currently limit this approach. However, the ongoing reduction in sequencing costs and the advanced discovery potential of long-read sequencing techniques point to a promising future in this research area. Despite the technological limitations in characterizing our dataset, we believe our analysis presents a valuable contribution to the Singapore and global scientific community. We advocate for the consensus *CYP2D6* star allele calling method used in our study for similar analyses, to address the challenges of short-read sequencing. Future studies focusing on the definitive characterization of the novel alleles not validated in this research, along with functional studies assessing their clinical significance, will be crucial for enhancing clinical pharmacogenomics implementation strategies. In the meantime, we believe that the detailed mapping of *CYP2D6* star allele distributions in Southeast Asian populations, as presented in our study, can serve as a resource in advancing precision medicine strategies and fostering the adoption of proactive pharmacogenetic testing in diverse clinical environments across Asia and worldwide.

## Methods

### Study population

Utilizing the SG10K_Health dataset^11^, our study evaluated 1,850 samples, each undergoing sequencing at a 30x depth. The ethnic composition showed a significant number of Chinese participants, comprising 74.6% (N=1,380) of the total. The Malay and Indian participants followed, contributing to 14.9% (N=275) and 10.5% (N=195) respectively. Participants’ ethnicities were determined based on their self-reported information.

### Consensus algorithm

Aldy^13^, Cyrius^12^, and StellarPGx^14^ are three well-established *CYP2D6* star allele callers designed for use with short-read data. We compared the star allele calls from these three tools to reach a consensus on the diplotypes assigned to each sample in our dataset, aiming to increase confidence in our callset.

The three tools were executed in parallel from CRAM inputs using a Nextflow workflow developed in-house. A custom parser read each sample’s output from Aldy, Cyrius, and StellarPGx. For ease of comparison across tools, parsed outputs were initially stored as separate haplotypes (e.g., *\*5/*36+*10* was split into *\*5* and *\*36+*10*). For callers that reported minor star alleles, these were excluded from the analysis since they are not functionally different from their corresponding major star alleles (e.g., *\*4C* was simplified to *\*4, *2.005* to *\*2*, and *\*2.ALDY* to *\*2*). Furthermore, the major star alleles of each haplotype were sorted in numerically ascending order and merged into diplotypes (e.g., *\*10+*36/*5* was rearranged to *\*5/*10+*36*) to facilitate downstream consensus comparison.

Processed diplotypes for each sample were then compared against each other. A consensus was considered achieved when more than 50% of calls supported a particular diplotype. Additionally, if the samples are found to have potential novel alleles, they were excluded from the mainstream consensus diplotype call-set and subjected to further manual inspections.

### Calculation of star allele frequencies and correlation with PharmGKB

The frequency of *CYP2D6* haplotypes was derived from the consensus diplotypes. We conducted a comparative analysis of the observed frequencies in the SG10K_Health dataset and those in PharmGKB^20^. For this comparison, we matched subsets of the population in each dataset, aligning East Asian with overall, Chinese, and Malay populations, and South Asian with the Indian population, as detailed in the PharmGKB *CYP2D6* frequency table (https://www.pharmgkb.org/page/cyp2d6RefMaterials). A notable limitation in our approach was the absence of frequency data for tandem variants (e.g., *\*36+*10* and *\*68+*4*) in the PharmGKB dataset. To address this, we supplemented our analysis with data from the 1000 Genomes Project (1KGP)^12^. Outliers in allele frequency between the two datasets were identified by calculating the frequency differences and considering a z-score greater than 3 as indicative of an outlier. Additionally, we assessed the correlation between the two datasets using Pearson’s correlation coefficient.

### Interpretation of star alleles into metabolizer profiles

Metabolizer profiles are categorized based on activity scores derived from the *CYP2D6* diplotypes. This scoring is based on the Clinical Pharmacogenetics Implementation Consortium (CPIC) guidelines^21^. The scoring system assigns function values to the star alleles (e.g., increased, normal, decreased, or no function). For each allele, an “activity value” ranging from 0 to 1 is assigned, such as 0 for no function, 0.5 for decreased function, and 1.0 for normal function. The activity score (AS) of a *CYP2D6* diplotype is then the sum of these values. Additionally, in cases where the *CYP2D6* allele has variable copy numbers, the activity value of an allele is multiplied by the number of gene copies. Within CPIC guidelines, metabolizer phenotypes are classified based on their total AS: individuals with an AS of 0 are poor metabolizers (PMs), those with a score between 0 and 1.25 are intermediate metabolizers (IMs), scores ranging from 1.25 −2.25 indicate normal metabolizers (NMs), and scores above 2.5 signify ultrarapid metabolizers (UMs).

### Identification and curation of novel alleles

The initial flagging of samples with potential novel alleles was conducted using StellarPGx. Overall, 93 samples were originally flagged and underwent a detailed investigation to infer the potential novel alleles. The initial step was seeking additional variants from the core variants reported by StellarPGx and ensuring the combination of variants or the additional variant itself were not already documented in PharmVar, suggesting a novel combination. Furthermore, the curation process involved manual variant inspection on the Integrative Genomic Browser (IGV) and verifying the number of reads supporting reference (REF) and alternative (ALT) alleles. To determine the background allele, effective phasing was achieved through two primary methods: either by identifying the same background allele present on both chromosomes, or by observing similar variant occurrences in multiple participants. Following this curation process, the original sample set was refined to 28 study samples containing 14 distinct novel alleles. For additional variants that are not documented in PharmGKB we inferred the variant classification using Variant Effect Predictor (VEP)^22^.

### Data availability

Data including WGS and intermediate files for all analyses and regeneration of all display items contain individual-level data including genotypes. The data is governed by the NPM Data Access Committee (DAC). The data generated in this study is made available to researchers registered through the SG10K_Health Data Access Portal. Users are required to submit a Data Access Request to the NPM DAC for approval. The forms and data access policy can be downloaded via the SG10K_Health web portal (https://npm.a-star.edu.sg/help/NPM). For more information, users can contact the National Precision Medicine Programme Coordinating Office, A*STAR.

### Grants

This study made use of data generated as part of the Singapore National Precision Medicine program funded by the Industry Alignment Fund (Pre-Positioning) (IAF-PP: H17/01/a0/007).

This study made use of data / samples collected in the following cohorts in Singapore:

- The Health for Life in Singapore (HELIOS) study at the Lee Kong Chian School of Medicine, Nanyang Technological University, Singapore (supported by grants from a Strategic Initiative at Lee Kong Chian School of Medicine, the Singapore Ministry of Health (MOH) under its Singapore Translational Research Investigator Award (NMRC/STaR/0028/2017) and the IAF-PP:H18/01/a0/016);
- The Growing up in Singapore Towards Healthy Outcomes (GUSTO) study, which is jointly hosted by the National University Hospital (NUH), KK Women’s and Children’s Hospital (KKH), the National University of Singapore (NUS) and the Singapore Institute for Clinical Sciences (SICS), Agency for Science Technology and Research (A*STAR) (supported by the Singapore National Research Foundation under its Translational and Clinical Research (TCR) Flagship Programme and administered by the Singapore Ministry of Health’s National Medical Research Council (NMRC), Singapore-NMRC/TCR/004-NUS/2008; NMRC/TCR/012-NUHS/2014. Additional funding is provided by SICS and IAF-PPH17/01/a0/005);
- The Singapore Epidemiology of Eye Diseases (SEED) cohort at Singapore Eye Research Institute (SERI) (supported by NMRC/CIRG/1417/2015; NMRC/CIRG/1488/2018; NMRC/OFLCG/004/2018);
- The Multi-Ethnic Cohort (MEC) cohort (supported by NMRC grant 0838/2004; BMRC grant 03/1/27/18/216; 05/1/21/19/425;11/1/21/19/678, Ministry of Health, Singapore, National University of Singapore and National University Health System, Singapore);
- The SingHealth Duke-NUS Institute of Precision Medicine (PRISM) cohort (supported by NMRC/CG/M006/2017_NHCS; NMRC/STaR/0011/2012, NMRC/STaR/ 0026/2015, Lee Foundation and Tanoto Foundation);
- The TTSH Personalised Medicine Normal Controls (TTSH) cohort funded (supported by NMRC/CG12AUG17 and CGAug16M012).

The views expressed are those of the author(s) are not necessarily those of the National Precision Medicine investigators, or institutional partners. We thank all investigators, staff members and study participants who made the National Precision Medicine Project possible.

