## Supplementary for "The variation landscape of *CYP2D6* in a multi-ethnic Asian population"

### Supplementary Materials

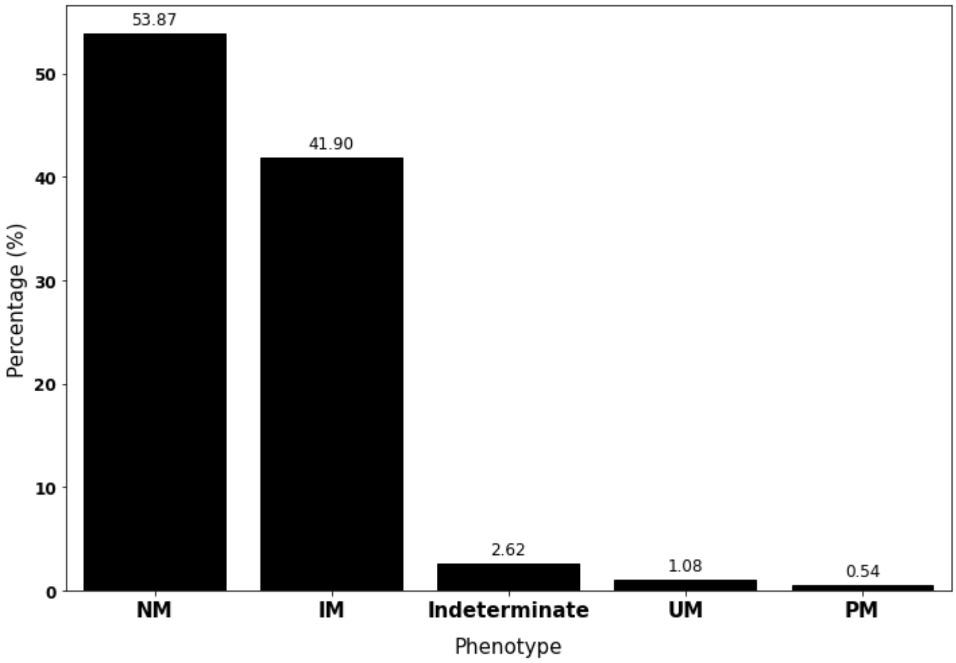


**Supplementary Figure 1: Frequency distribution of CYP2D6 phenotypes (N=1487).** The x-axis lists all the possible CYP2D6 phenotypes, including normal metabolizers (NM), intermediate metabolizers (IM), indeterminate, ultrarapid metabolizers (UM), and poor metabolizers (PM). The y-axis shows the frequency distribution (%) of these phenotypes across each genetic ancestry.

**Supplementary Table 1: Frequencies of *CYP2D6* star alleles in the SG10K_Health dataset, categorized by genetic ancestries.**

| | | Frequency (%) | | | |
|---|---|---|---|---|---|
| **Star Allele** | **CPIC Function** | **Overall (N=2974)** | **Chinese (N=2248)** | **Indian (N=328)** | **Malay (N=398)** |
| **1* | Normal function | 27.81 | 25.85 | 37.80 | 30.65 |
| **2* | Normal function | 9.92 | 8.59 | 19.51 | 9.55 |
| **35* | Normal function | 0.50 | 0.44 | 1.52 | 0 |
| **39* | Normal function | 0.13 | 0.13 | 0 | 0.25 |
| **33* | Normal function | 0.07 | 0 | 0.61 | 0 |
| **2x2* | Increased function | 0.34 | 0.31 | 0.30 | 0.50 |
| **1x2* | Increased function | 0.30 | 0.09 | 1.22 | 0.75 |
| **10+*36* | Decreased function | 28.95 | 34.30 | 3.05 | 20.10 |
| **10* | Decreased function | 17.45 | 18.02 | 3.35 | 25.88 |
| **41* | Decreased function | 4.74 | 3.74 | 12.20 | 4.27 |
| **14* | Decreased function | 1.08 | 1.38 | 0 | 0.25 |
| **49* | Decreased function | 0.71 | 0.89 | 0 | 0.25 |
| **17* | Decreased function | 0.03 | 0 | 0.30 | 0 |
| **9* | Decreased function | 0.03 | 0 | 0.30 | 0 |
| **5* | No function | 3.43 | 3.16 | 6.10 | 2.76 |
| **36* | No function | 1.58 | 1.87 | 0 | 1.26 |
| **4* | No function | 1.01 | 0.04 | 6.40 | 2.01 |
| **4+*68* | No function | 0.34 | 0.04 | 2.74 | 0 |
| **21* | No function | 0.13 | 0.18 | 0 | 0 |
| **7* | No function | 0.03 | 0 | 0.30 | 0 |
| **15* | No function | 0.03 | 0 | 0 | 0.25 |
| **4+*4* | No function | 0.03 | 0 | 0.30 | 0 |
| **69* | No function | 0.03 | 0.04 | 0 | 0 |
| **71* | Uncertain function | 0.44 | 0.44 | 0 | 0.75 |
| **86* | Uncertain function | 0.20 | 0.09 | 1.22 | 0 |
| **43* | Uncertain function | 0.20 | 0.09 | 1.22 | 0 |
| **94* | Uncertain function | 0.10 | 0.13 | 0 | 0 |
| **65* | Uncertain function | 0.07 | 0.04 | 0 | 0.25 |
| **113* | Uncertain function | 0.07 | 0 | 0.61 | 0 |
| **111* | Uncertain function | 0.07 | 0 | 0.30 | 0.25 |
| **112* | Uncertain function | 0.03 | 0 | 0.30 | 0 |
| **52* | Uncertain function | 0.03 | 0.04 | 0 | 0 |
| **133* | Uncertain function | 0.03 | 0.04 | 0 | 0 |
| **75* | Uncertain function | 0.03 | 0.04 | 0 | 0 |
| **82* | Uncertain function | 0.03 | 0 | 0.30 | 0 |

CPIC, Clinical Pharmacogenetics Implementation Consortium.
